# Diversity and determinants of recombination landscapes in flowering plants

**DOI:** 10.1101/2022.03.10.483889

**Authors:** Thomas Brazier, Sylvain Glémin

## Abstract

During meiosis, crossover rates are not randomly distributed along the chromosome and therefore they locally influence the creation of novel genotypes and the efficacy of selection. To date, the broad diversity of recombination landscapes among plants has rarely been investigated, undermining the overall understanding of the constraints driving the evolution of crossover frequency and distribution. The determinants that shape the local crossover rate and the diversity of the resulting landscapes among species and chromosomes still need to be assessed in a formal comparative genomic approach. We gathered genetic maps and genomes for 57 flowering plant species, corresponding to 665 chromosomes, for which we estimated large-scale recombination landscapes. Chromosome length drives the basal recombination rate for each species, but within species we were intrigued to notice that the chromosome-wide recombination rate is proportional to the relative size of the chromosome. Moreover, for larger chromosomes, crossovers tend to accumulate at the ends of the chromosome leaving the central regions as recombination-free regions. Based on identified crossover patterns and testable predictions, we proposed a conceptual model explaining the broad-scale distribution of crossovers where both telomeres and centromeres are important. Finally, we qualitatively identified two recurrent crossover patterns among species and highlighted that these patterns globally correspond to the underlying gene distribution. In addition to the positive correlation between recombination and gene density, we argue that crossover patterns are essential for the efficiency of chromosomal genetic shuffling, even though the ultimate evolutionary potential forged by the diversity of recombination landscapes remains an open question.

## Introduction

Recombination is a universal feature of sexually reproducing species that participates to the production of new haplotypes passed on to offspring by the reciprocal exchange of DNA between maternal and paternal chromosomes during meiosis. However, recombination landscapes are not homogeneous along the chromosome and vary among species (de Massy, 2013; Haenel et al., 2018; Mézard et al., 2015; Stapley et al., 2017). Meiotic recombination plays a fundamental functional role by forming chiasmata at specific pairing sites between homologous chromosomes to ensure the physical tension needed for the proper disjunction of homologs (de Massy, 2013; Mézard et al., 2015; Zickler and Kleckner, 2015). Recombination also plays an evolutionary role by breaking the linkage between neighbouring sites and creating new genetic combinations transmitted to the next generation upon which selection can act (Barton, 1995; Charlesworth and Jensen, 2021; Otto, 2009). However, the number and location of crossovers (COs) along the chromosome are finely regulated through mechanisms of crossover assurance, interference and homeostasis (Otto and Payseur, 2019; Pazhayam et al., 2021). In most species, at least one CO per chromosome is mandatory to achieve proper segregation and to avoid deleterious consequences of nondisjunction. Additional COs are also usually regulated through interference, ensuring that they are not too numerous and not too close to each other (Pazhayam et al., 2021; Wang et al., 2015). In addition to regulation on a large scale (Cooper et al., 2016; Zelkowski et al., 2019), recombination is also finely tuned on a small scale. Generally, crossovers are concentrated in very short genomic regions (typically a few kb), i.e. recombination hotspots. In plants, CO hotspots are usually found in gene regulatory sequences, and mostly in promoters (Choi et al., 2018; He et al., 2017; Marand et al., 2019).

In addition to meiosis functioning, variations in recombination rates have a strong impact on genome structure, functioning and evolution (Gaut et al., 2007; Haenel et al., 2018; Stapley et al., 2017; Tiley and Burleigh, 2015). For example, recombination landscapes are thought to shape genetic diversity and the distribution of transposable elements (TEs) along chromosomes, through the indirect effect of recombination in modulating the extent of linked selection (Corbett-Detig et al., 2015; Kent et al., 2017). Recombination can also shape nucleotide landscapes through the effect of GC-biased gene conversion (Galtier et al., 2018; Glémin et al., 2014). As a consequence, it has become a key challenge to integrate recombination rate variation in population genomics in the age of ‘genomic landscapes’ (Booker et al., 2020; Comeron, 2017). The characterization of recombination landscapes also has practical interests since it is likely that changes in CO patterns is to be manipulated for an advantageous purpose, such as creating specific genetic combinations or directly influencing genetic diversity and the adaptive potential of a species (Kuo et al., 2021).

To date, the diversity of recombination landscapes in plants has not been properly quantified; it is often limited to genome-wide recombination rates (Stapley et al., 2017), even though it could be used as a lever to identify the major determinants shaping crossover patterns across species (Gaut et al., 2007). Recently, a meta-analysis explored large-scale recombination landscapes among eukaryotes and concluded that chromosome length plays a major role in crossover patterns, but this analysis only included a limited number of plant species (Haenel et al., 2018). As plant genomes are highly diverse in many ways (Pellicer et al., 2018; Soltis et al., 2015), the expected diversity in recombination landscapes has been overlooked (Gaut et al., 2007). Plant genomes contain large regions with suppressed recombination, which impact genomic averages (Gaut et al., 2007; Haenel et al., 2018), and it seems that the physical location of COs along the chromosome is species-specific (Wang and Copenhaver, 2018). Nonetheless, several broad-scale determinants have recently been identified, such as chromosome length (Haenel et al., 2018; Tiley and Burleigh, 2015), distance to the telomere or centromere (Blitzblau et al., 2007; Haenel et al., 2018) and genomic and epigenetic features (Apuli et al., 2020; Marand et al., 2019; Yelina et al., 2012). In plants, recombination rates are supposed to be higher in smaller genomes because the linkage map length is independent of genome size (Stapley et al., 2017) and the existence of a major broad-scale determinant such as chromosome size needs to be tested. Haenel et al. (2018) suggested a simple telomere-led model with a universal bias of COs towards the periphery of the chromosome, driven by chromosome length; this new conceptual model still needs to be tested on data. However, there is little evidence supporting a universal pattern among plants (Zelkowski et al., 2019) and it has been proposed that both telomeres and centromeres shape recombination landscapes, although this is not yet fully understood (Wang and Copenhaver, 2018). Since recombination hotspots are supposedly found in gene regulatory sequences, gene density could also be a universal driver of recombination rates among plant species, but this still needs to be tested.

By combining genetic mapping from pedigree data and genome assembly up to the chromosome scale, we have gathered, to the best of our knowledge, the largest recombination landscape dataset in flowering plants. More precisely, we have estimated the rate of COs along chromosomes. We tested the relationship between recombination and chromosome length and assessed if the distribution of COs could be shaped by genome structure (i.e. chromosome size, telomeres, centromeres). We identified two main patterns of recombination that are parallel to, and which may emerge from, the gene density distribution. Finally, we discussed the possible evolutionary implications of the heterogeneity of recombination landscapes by quantifying how CO patterns affect genetic shuffling, which raises new questions on the evolution of recombination.

## Results

### Dataset and recombination maps

We retrieved publicly available data for linkage maps and genome assemblies. We selected linkage maps for which the markers had genomic positions on a chromosome-level genome assembly (except for *Capsella rubella*, which had a high-quality scaffold-level assembly, i.e. pseudo-chromosomes). After making a selection based on the number of markers, marker density, and genome coverage, and after filtering out the outlying markers, we adopted a qualitative visual validation. In the end, we retained 665 chromosome-scale Marey maps (plot of the genetic vs genomic distance, cM vs Mb) for 57 species (2-26 chromosomes per species), from which we successfully inferred recombination landscapes (Table S1, S2, Figures S1, S2). After removing the outliers, the number of markers per chromosome map ranged from 31 to 49,483, with a mean of 956 markers per map. We used a linear regression on the models’ residuals to verify that neither the number of markers, marker density nor the number of progenies had a significant effect on the analyses. We also retrieved gene annotations for 41 genomes. The angiosperm phylogeny was well represented in our sampling (Figure S3), with a basal angiosperm species (*Nelumbo nucifera*), 15 monocot species and 41 eudicots. We also searched the literature for data on the centromeric index for 37 species, defined as the ratio of the short arm length divided by the total chromosome length (Table S3).

From the Marey maps, we estimated local recombination rates along the chromosomes on non-overlapping 100 kb windows. Estimates at a scale of 1 Mb yielded very similar results (correlation between 1 Mb windows and 100 kb windows pooled in 1 Mb windows, Spearman rank correlation coefficient Rho = 0.99, p < 0.001, Table S4) therefore only 100 kb landscapes were analysed in the subsequent analyses.

### Smaller chromosomes recombine more often than larger ones

Our results are in agreement with previous studies showing that smaller chromosomes have a higher recombination rate per Mb than larger ones (Haenel et al., 2018; Stapley et al., 2017), and our sampling suggests a consistent pattern across species (Figure 1A). We found a significant negative correlation between chromosome size (Mb) and the mean chromosomal recombination rate (Spearman rank correlation coefficient Rho = −0.84, p < 0.001; log-log Linear Model, adjusted R^2^ = 0.83, p < 0.001). For most species, there were between one and four COs per chromosome, which suggests that the number of COs per chromosome remains stable across species even though the genome sizes span almost two orders of magnitude.

**Figure 1.**
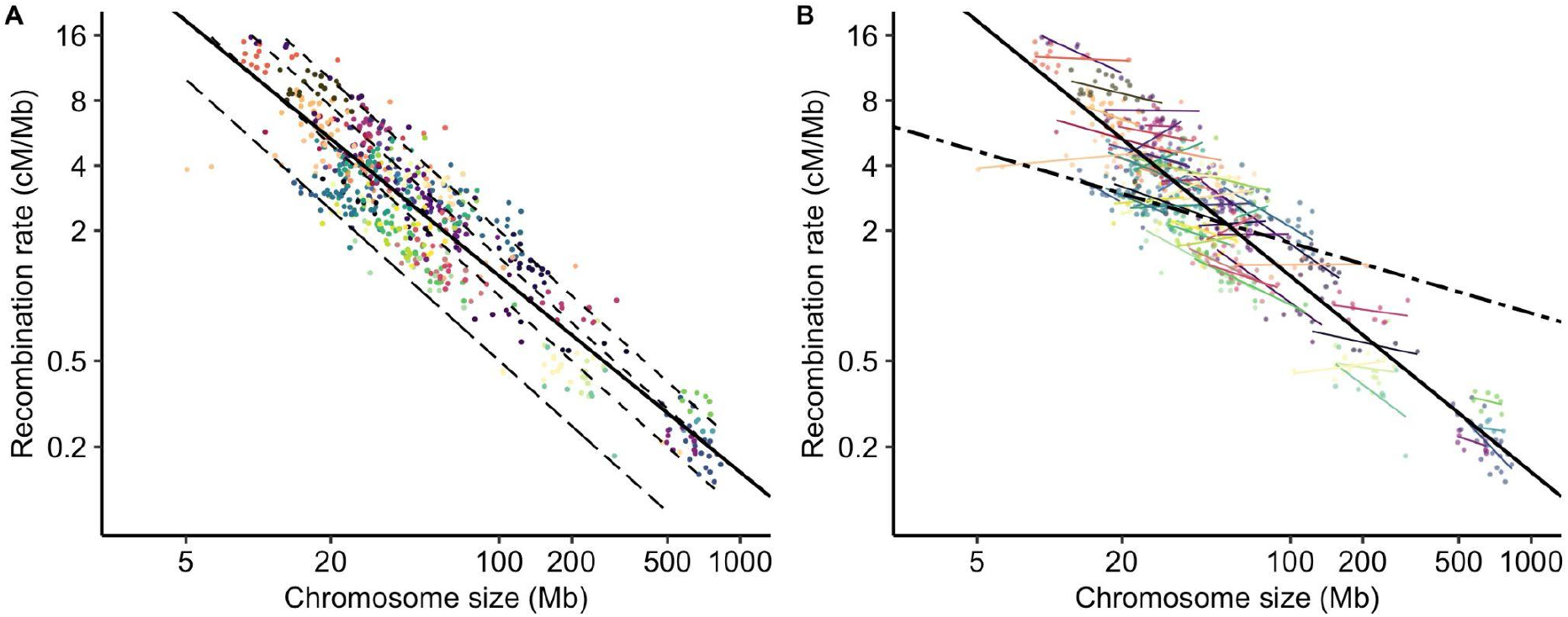
Recombination rates (cM/Mb, log scale) are negatively correlated to chromosome genomic size (Mb, log scale). Recombination rates were estimated with the loess regression function in windows of 100 kb and averaged per chromosome. Each point represents a chromosome (n = 665). Species are presented in different colours (57 species). (A) The black solid line represents the Linear Model regression line fitted to the data (log-log Linear Model, adjusted R^2^ = 0.83, p < 0.001). The lower long dashed line corresponds to the expectation of one CO per chromosome, and the upper dashed lines correspond to two, three or four COs respectively (ascending order). (B) Correlations between recombination rates and chromosome size within species and a between-species correlation controlled for a species effect (n = 57 species). The black dashed line represents the selected Linear Mixed Model with a species effect (log-log LMER, marginal R^2^ = 0.17, conditional R^2^ = 0.96, p < 0.001). Coloured lines show the random regressions for species (55 species regression lines for species with at least 5 chromosomes, 5-26 chromosomes per species).

Using a Linear Mixed Model (LMER) we found a significant species random effect for both the intercept and the slopes (log-log LMER, marginal R^2^ = 0.17, conditional R^2^ = 0.96, p < 0.001); the introduction of phylogenetic covariance did not improve the mixed model (Table S5). Interestingly, the LMER results showed that the (log-log) relationship between the recombination rate and the chromosome size was not the same within and between species, suggesting that absolute chromosome size does not have a general effect among species (Figure 1B). Similarly, the relationship between linkage map length (cM) and chromosome size (Mb) was highly species specific (log-log linear mixed model, marginal R2 = 0.49, conditional R2 = 0.99, p < 0.001) (Figure 2), with species slopes decreasing with the mean chromosome size in a log-log relationship, indicating that species slopes are roughly proportional to the inverse of the mean chromosome size (Figure 2C). As a consequence, the excess of COs on a chromosome (i.e. the linkage map length minus 50 cM) was not correlated to the absolute chromosome size but to the relative one (i.e. chromosome size divided by the mean chromosome size of the species). Moreover, in contrast to the relationship between recombination rate and absolute size, we did not observe any difference between the linear model and the fixed regression of the mixed linear model, suggesting that this relationship is similar across species (Figure 2D).

**Figure 2.**
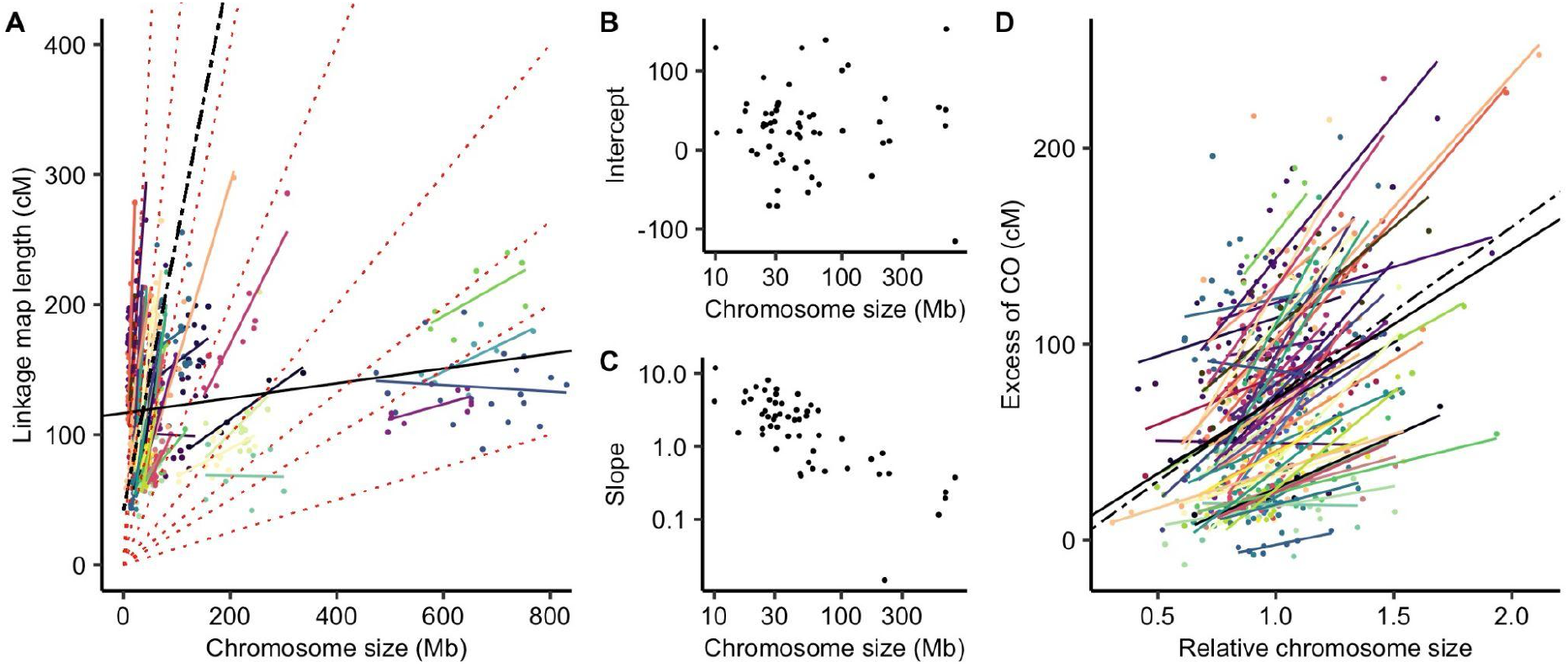
Linkage map length (cM) is positively correlated to genomic chromosome size (Mb). (A) Correlation between chromosome genomic size (Mb) and linkage map length (cM). Each point represents a chromosome (n = 665). Species are presented in different colours (57 species). The linear regression is the solid black line (Linear Model, adjusted R^2^ = 0.036, p < 0.001). The fixed regression of the Linear Mixed Model is the black dashed line (LMER, marginal R^2^ = 0.49, conditional R^2^ = 0.99, p < 0.001). Species random slopes are shown in colour. Isolines of the Genome-wide Recombination Rate (GwRR) were plotted as dotted red lines to represent regions with equal recombination rates. (B) Random intercepts for species as a function of the species mean genomic chromosome size (Mb). (C) Random slopes for species as a function of the species mean genomic chromosome size (Mb). (D) The excess of COs (i.e. linkage map length minus 50 cM for the obligate CO) is consistently positively correlated to the relative chromosome size (i.e. chromosome size divided by the averaged chromosome size of the species). Each point represents a chromosome (n = 665). Species are presented in different colours (57 species). The black solid line is the linear regression across species (Linear Model, adjusted R^2^ = 0.13, p < 0.001). The black dashed line is the linear mixed regression with a random species effect (LMER, marginal R^2^ = 0.14, conditional R^2^ = 0.86, p < 0.001). Coloured solid lines represent individual regression lines (Linear Model) for species with at least 5 chromosomes (55 species, 5-26 chromosomes per species).

### Diversity of CO patterns among flowering plants

Recombination landscapes along chromosomes appeared to be qualitatively very similar within species but strongly varied between species (Figure 3, Figure S2). COs were not evenly distributed between the centre and extremities of the chromosomes. In the text below, we have used the terms proximal and distal regions, respectively, to avoid confusion with the molecular composition and specific position defining telomeric and centromeric regions *stricto sensu*. Some landscapes were homogeneous along chromosomes whereas others were extremely structured with the recombination concentrated in the short distal parts of the genome, with wide pattern variations between these two extremes (Figure 3). Using a broken stick representation where relative recombination rates were presented on ten bins of equal chromosome length (see Materials and Methods for details), we observed that the bias towards the periphery was not ubiquitous across species (Figure 4); Haenel et al. (2018) concluded to a distal bias universal for chromosomes larger than 30 Mb. Only a subset of species, especially those with larger chromosomes, exhibited a clear bias, with COs clustered in the distal regions and with recombination rates that were lower than expected at the centre (Figure 4). Despite large chromosome sizes (mean chromosome sizes = 101 Mb and 198 Mb, respectively), *Nelumbo nucifera* and *Camellia sinensis* are noticeable exceptions to this pattern, with the highest recombination rates found in the middle of the chromosomes (*Nelumbo nucifera* illustrated in Figure 3E, other species in Figure S2). For small to medium-sized chromosomes, the pattern is less clear. Most species did not show any clear structure along the chromosome but a few of them (e.g. *Capsella rubella, Dioscorea alata, Mangifera indica, Manihot esculenta)* showed a drop in recombination rates in the distal regions and high recombination rates in the proximal regions (*Capsella rubella* illustrated in Figure 3A).

**Figure 3.**
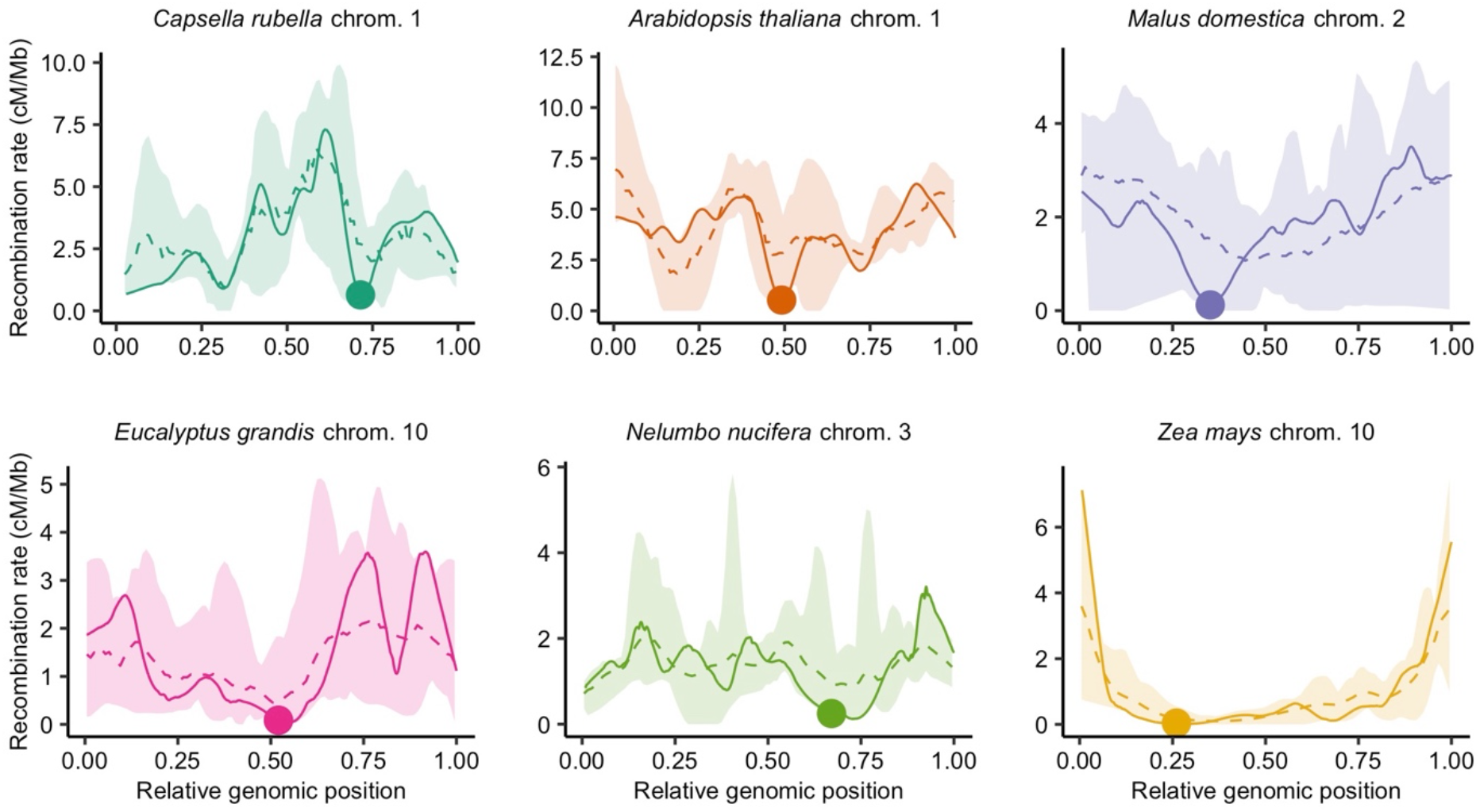
The diversity of recombination landscapes in angiosperms is exemplified by six different emblematic species. Recombination landscapes are similar within species (the dashed line is the average landscape for pooled chromosomes, all recombination landscapes of the species are contained within the colour ribbon). Genomic distances (Mb) were scaled between 0 and 1 (divided by chromosome size) to compare chromosomes with different sizes. Estimates of the recombination rates were obtained by 1,000 bootstrap replicates of loci in windows of 100 kb with loess regression and automatic span calibration. The averaged species recombination landscape (dashed line) was estimated by calculating the mean recombination rate in 100 bins along the chromosome axis, all chromosomes pooled. Similarly, the lower and upper boundaries (pale ribbon) were estimated by taking the minimum and maximum recombination rates in 100 bins. One chromosome per species is represented in a solid line, with the genomic position of the centromere demarcated by a dot. The six species are ordered by ascending mean chromosome size (Mb).

**Figure 4.**
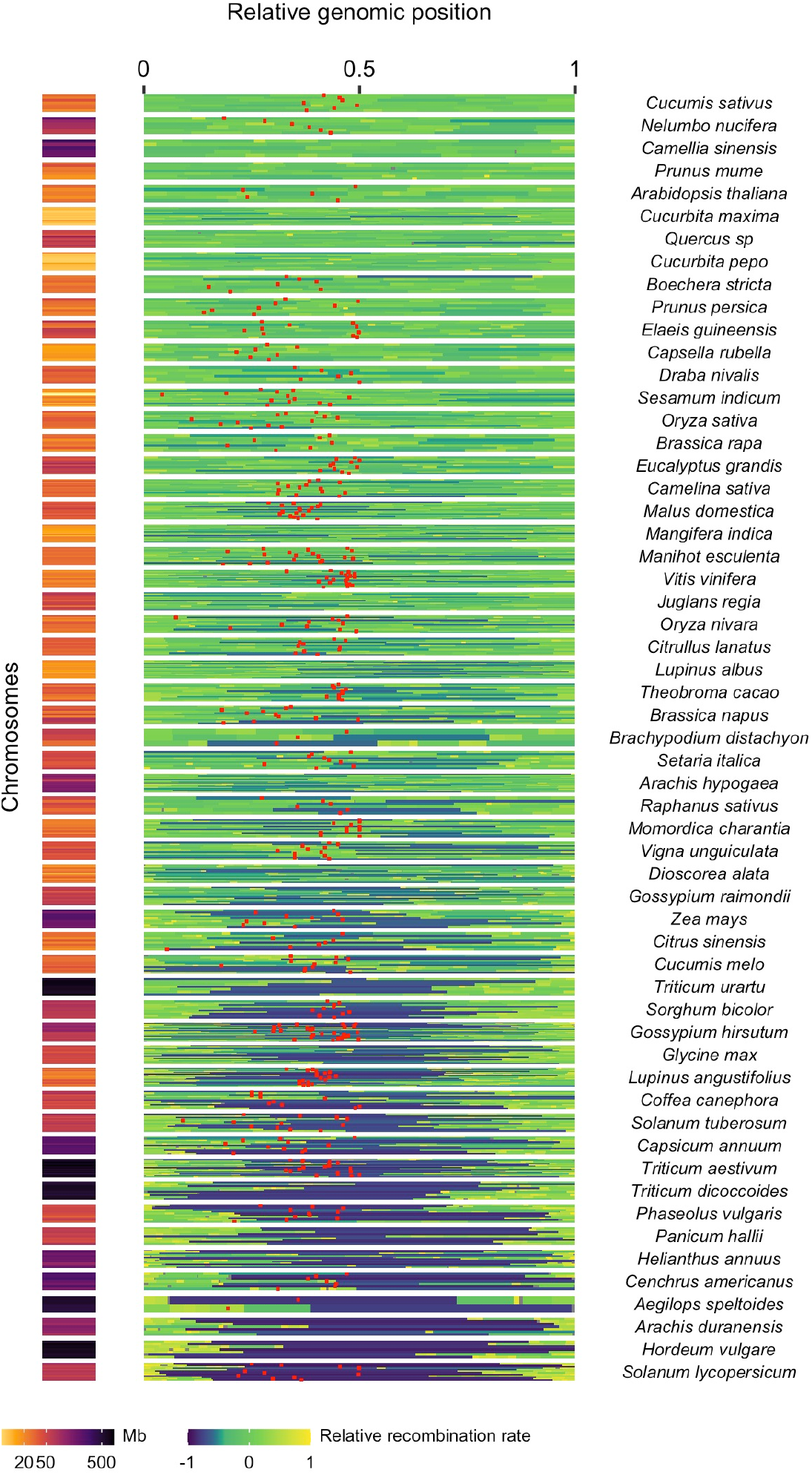
Patterns of recombination within chromosomes (n = 665). Relative recombination rates along the chromosome were estimated in ten bins using the broken stick model. Species are ordered by ascending (top to bottom) variation in the relative recombination rates (57 species). Each horizontal bar plot represents the spatial distribution of recombination along a chromosome. Each chromosome was divided in ten bins of equal genomic size, i.e. 1/10 of the total genomic size (Mb). The relative recombination rate is the log-transformed ratio of the expected relative genetic length (one tenth) divided by the observed relative genetic length of the bin. It means that values below zero are recombination rates that are lower than expected under a random distribution of COs whereas values above zero are recombination rates that are higher than expected. Whenever the centromere position on the chromosome is available, this information is mapped as a red dot and chromosomes are oriented with the longer arm on the right. Chromosome sizes (Mb) on the left correspond to each broken stick chromosome.

Following Haenel et al. (2018), we calculated the periphery-bias ratio as the recombination rate in the tips of the chromosome (10% at each extremity) divided by the mean recombination rate. A ratio higher than 1 indicates a higher recombination rate in the tips than the whole chromosome. By pooling chromosomes per species, we detected a significant positive effect of chromosome length on the periphery-bias ratio across species (Spearman rank correlation coefficient Rho = 0.60, p < 0.001; Linear Model, adjusted R^2^ = 0.44, p < 0.001) (Figure 5A). At the chromosome level, the mean periphery-bias ratio is significantly higher than 1 (95% bootstrapped confidence interval = [2.06;2.32]) but the correlation with chromosome length within species was not clear (Figure 5B, 5C, Table S6). Although we do find some ratios below 1 (Figure 5B), the distribution of the periphery-bias ratios is clearly skewed towards values higher than 1, suggesting that spatial clustering in the tips of the chromosome is a common feature among angiosperms, however with many exceptions (Figure 3A, 3E).

**Figure 5.**
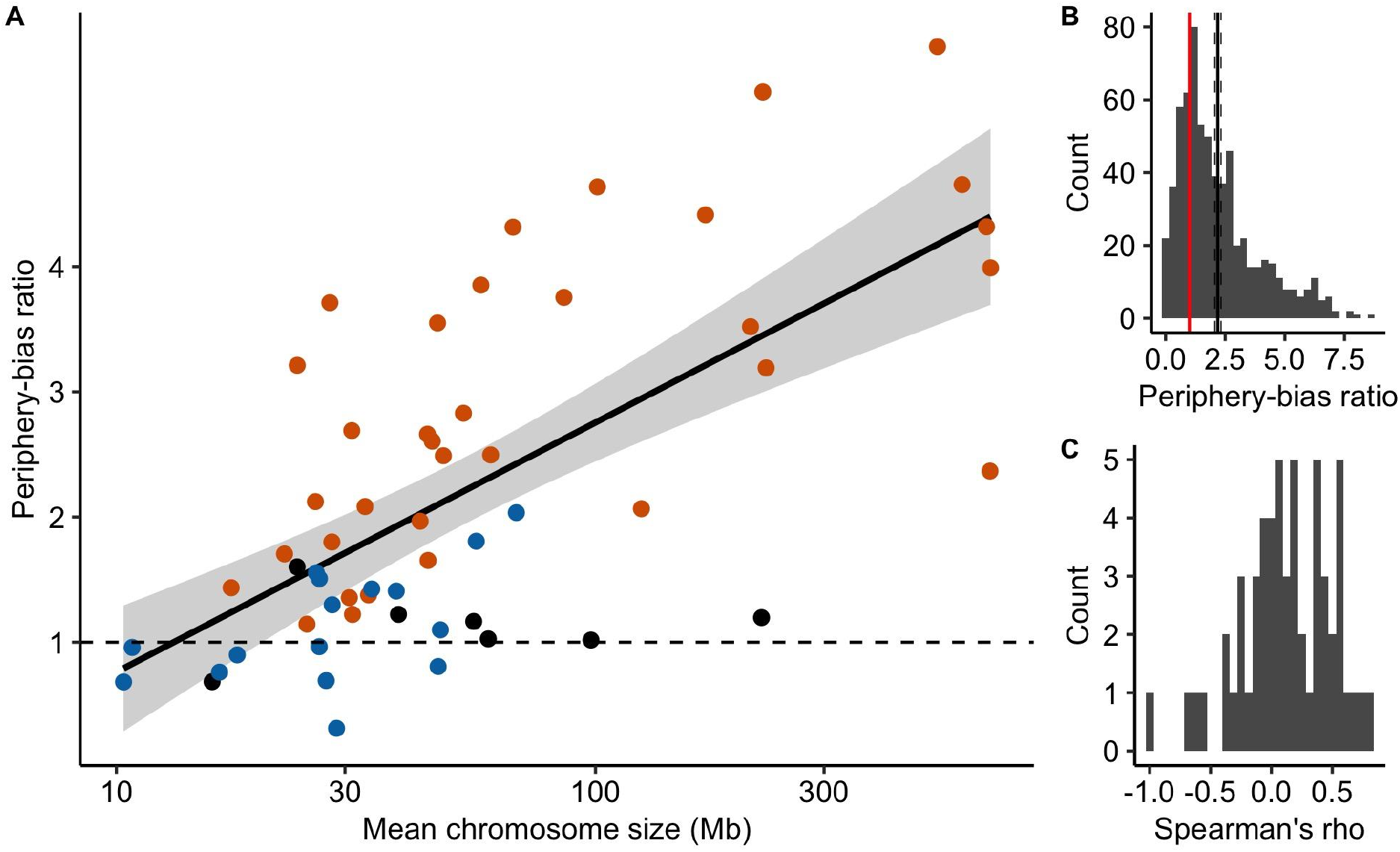
The periphery-bias ratio is positively correlated to chromosome genomic size. (A) The periphery-bias ratio depends on the mean chromosome size (Mb, log scale) across species (n = 57 species). The linear regression line and its parametric 95% confidence interval were estimated in ggplot2 (Linear Model, chromosome size log-transformed, adjusted R^2^ = 0.44, p < 0.001). A periphery-bias ratio above 1 (dashed horizontal line) indicates that recombination rates in the tips of the chromosome are higher than the mean chromosome recombination rate. Points are coloured according to the classification of the CO patterns described below (orange = distal, blue = sub-distal, black = unclassified). (B) Distribution of periphery-bias ratios (n = 665 chromosomes). The mean periphery-bias ratio and its 95% confidence interval (black solid and dashed lines) were estimated by 1,000 bootstrap replicates. The red vertical line shows the theoretical value for an equal recombination in the tips compared to the rest of the chromosome (periphery-bias ratio = 1). (C) Distribution of Spearman’s correlation coefficients between the periphery-bias ratio and chromosome genomic size (Mb) within species (n = 57 species).

### Joint effect of telomeres and centromeres on crossover distribution along chromosomes

Globally, recombination rates were negatively correlated to the distance to the nearest telomere (Figure S4, Table S7, Table S8). However, two different patterns qualitatively emerged (Figure 6, Figure S5, Table S8). In 34 species, recombination decreased from the telomere and reached a plateau after a relative genomic distance of approximately 20% of the whole chromosome (the distal model), in agreement with the model suggested by Haenel et al. (2018). Sixteen species presented a sharp decrease in the most distal regions and a peak of recombination in the sub-distal regions (relative genomic distance between 0.1-0.2) followed by a slow decrease towards the centre of the chromosome (the sub-distal pattern). There were very few exceptions to these two patterns (six species), e.g. *Capsella rubella* consistently showed higher recombination rates in the middle of the chromosome (Figure 3A). Interestingly, chromosomes from species classified as having a distal pattern were significantly larger than chromosomes with a sub-distal pattern (Wilcox rank sum test, p < 0.001, Figure 6C). Furthermore, the species correlation was significantly negatively correlated to the mean chromosome length (Spearman rank correlation coefficient Rho = - 0.51, p < 0.001; Figure S4).

**Figure 6.**
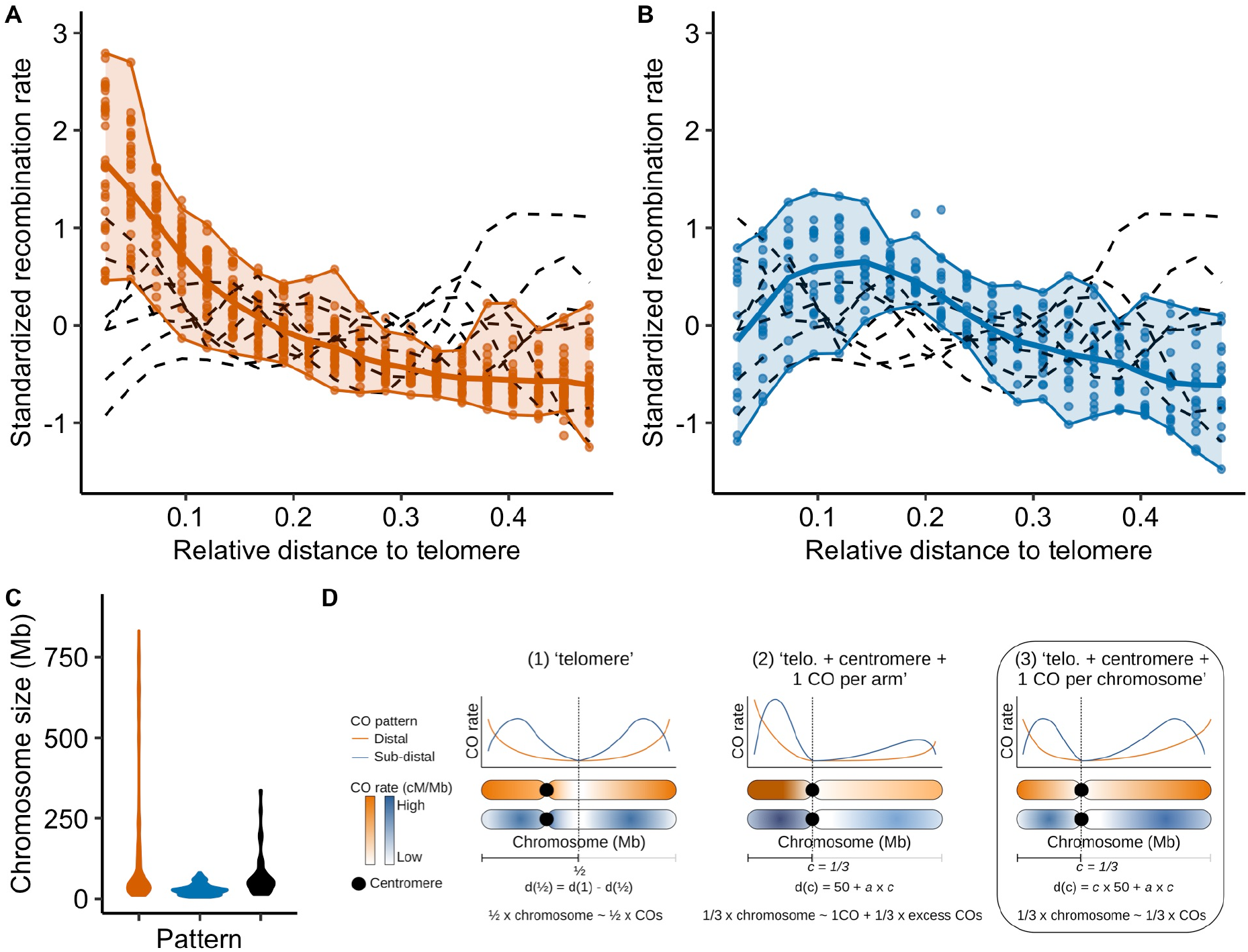
Crossover patterns can be classified in two different patterns, where recombination rates are higher in distal regions and lower near the centre of the chromosome. Standardized recombination rates for species (centred-reduced cM/Mb, chromosomes pooled per species, n = 57 species) are expressed as a function of the relative genomic distance from the telomere in 20 bins. Two patterns were identified and species were pooled accordingly, with 7 unclassified species (orange = distal, blue = sub-distal, black = unclassified). (A) In the distal pattern (34 species), recombination rates decreased immediately from the tip of the chromosome (left plot, orange line and ribbon). (B) In the sub-distal pattern (16 species), recombination rates were reduced in the distal regions and the peak of recombination was in the sub-distal region (right plot, blue line and ribbon). In each plot, the solid line represents the mean recombination rate estimated in a bin (20 bins). Chromosomes were split in half, where a distance of 0.5 is the centre of the chromosome. Then, chromosomes were pooled per species (each point is the mean recombination rate of all chromosomes in a species, for a distance bin to the tip of the chromosome). Upper and lower boundaries of the ribbon represent the maximum and minimum values attained for a particular pattern. Patterns that were not classified (7 species) were represented by individual loess regressions using black dashed lines. (C) Distribution of chromosome genomic sizes (Mb) in each pattern. (D) Three competing models compatible with the distal and sub-distal crossover patterns were tested using formal tests on predictions. Models are compatible with the distal and sub-distal crossover patterns we identified. Predictions are made on the distribution of crossovers between both chromosome sides (separated by the middle of the chromosome, model 1) or chromosome arms (separated by the centromere, models 2 and 3). Symbolic chromosomes illustrate the distribution of crossovers expected under each model and for each pattern. Model 3 is the best model (box).

When the centromere position was known, we qualitatively observed that the centromeres had an almost universal local suppressor effect (Figure 3). In small and medium-sized chromosomes, the recombination was often suppressed in short restricted centromeric regions (several Mb) displaying drastic drops in the recombination rates, whereas the rest of the map did not seem to be affected. In larger chromosomes, the suppression of recombination extends to large regions upstream and downstream of the physical centre of the chromosome (approximately 80-90% of the chromosome; Figure 4).

To go further, we formally compared three models that could explain the broad-scale crossover patterns we observed (Figure 6D). Under the strict distal model proposed by Haenel et al. (2018), the centromere does not play any role beyond its local suppressor effect, and therefore an equal distribution of crossovers on both sides of the chromosome is expected, independently of centromere position. In other words, we should expect 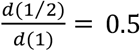, where *d*(1/2) is the genetic distance (cM) to the middle of the chromosome and *d*(1) is the total genetic distance (cM). In addition to this model (M1), we tested two nested models adding a centromere effect. We assumed that the position of the centromere, *d*(*c*), has an effect on the distribution of crossovers along the chromosome. Model M2 ‘telomere + centromere + one CO per arm’ and M3 ‘telomere + centromere + one CO per chromosome’ assume that the relative genetic distance of a chromosome arm is proportional to its relative genomic size. However, models M2 and M3 differ in the number and distribution of mandatory COs. At least one CO in each chromosome arm (50 cM) is mandatory in M2 whereas only one CO is mandatory for the entire chromosome in M3. Based on goodness-of-fit criteria (adjusted R^2^, AIC, BIC), we used linear regression to compare the theoretical predictions of the three competing models to the observed values for Marey maps in which the centromere position was known (37 species, 425 chromosomes). Model M2 was generally rejected since 22% of chromosomes showed less than 50 cM in at least one arm, even though it was supported in a handful of species (Table 1), and model M1 was not supported by any species. Model M3 was the best supported model (30 out of 37 species), with good predictive power (Spearman rank correlation between predicted and observed values: Rho = 0.72, p < 0.001; Tables 1, S9, S10). Given that some chromosomes had one chromosome arm shorter than 50 cM, which is incompatible with one mandatory CO per arm in model M2, we performed a second model selection on a subset of chromosomes with at least 50 cM on each chromosome arm (n = 36 species, 333 chromosomes) which confirmed that model M3 was the best model.

**Table 1.** Model selection for the telomere/centromere effect (n = 37 species with a centromere position, 425 chromosomes). Three competing models were compared based on the adjusted R^2^ p-value and AIC-BIC criteria among chromosomes (the best supported model is in bold characters). The number of species supporting each model was calculated based on the adjusted R^2^ within species, for all species with at least five chromosomes. (1) ‘telomere’ model. (2) ‘telomere + centromere + one CO per arm’ model. (3) ‘telomere + centromere + one CO per chromosome’ model. *d*(*c*) is the genetic distance to the centromere. *d*(1) is the total genetic distance. A second model selection was done on a subset of chromosomes with at least 50 cM on each chromosome arm (n = 36 species, 333 chromosomes).

| # | Model | Expected | Adjusted $R^2$ | p | AIC | BIC | Species |
| --- | --- | --- | --- | --- | --- | --- | --- |
| Full dataset (37 species, 425 chromosomes) |  |  |  |  |  |  |  |
| 1 | Telomere | $d(1/2) / d(1) = 0.5$ | 0.22 | < 0.001 | -477.8 | -465.7 | 0 |
| 2 | Tel. + Cent. + CO per arm | $(d(c) - 50) / (d(1) - 100) = c$ | - | 0.72 | 3098.2 | 3110.4 | 7 |
| 3 | Tel. + Cent. + CO per chr. | $d(c) / d(1) = c$ | 0.51 | < 0.001 | -476.6 | -464.5 | 30 |
| Subset (36 species, 333 chromosomes) |  |  |  |  |  |  |  |
| 1 | Telomere | $d(1/2) / d(1) = 0.5$ | 0.18 | < 0.001 | -407.5 | -396.1 | 0 |
| 2 | Tel. + Cent. + CO per arm | $(d(c) - 50) / (d(1) - 100) = c$ | -0.001 | 0.42 | 1939.1 | 1950.5 | 10 |
| 3 | Tel. + Cent. + CO per chr. | $d(c) / d(1) = c$ | 0.50 | < 0.001 | -396 | -384.6 | 26 |

### Recombination rates are positively correlated to gene density

At a fine scale, it has been shown in a few species that COs preferentially occur in gene promoters. The scale of 100 kb used here is too large to directly test whether this is a common pattern shared among angiosperms. Instead, we assessed whether recombination increased with gene density. Forty-one genomes were annotated with gene positions. Across chromosomes, the distribution of chromosomal correlations between gene count and recombination rate was clearly skewed towards positive values, independently of the previously described CO patterns (mean Spearman’s rank correlation = 0.46 [0.43; 0.49]; Figure 7A). Ninety-one percent (91%) of 483 chromosomes (41 species) showed a significant correlation between the number of genes and recombination rate at a 100 kb scale. Yet the strength of the relationship greatly varied across species and did not correlate with usual predictors such as the chromosome length or the genome-wide recombination rate (Figure 7B). Overall, standardized recombination rates (subtracting the mean and dividing by the standard deviation to allow comparison among species) consistently increased with the number of genes in most species (linear quadratic regression, adjusted R2 = 0.62, p < 0.001; Figure 7C).

**Figure 7.**
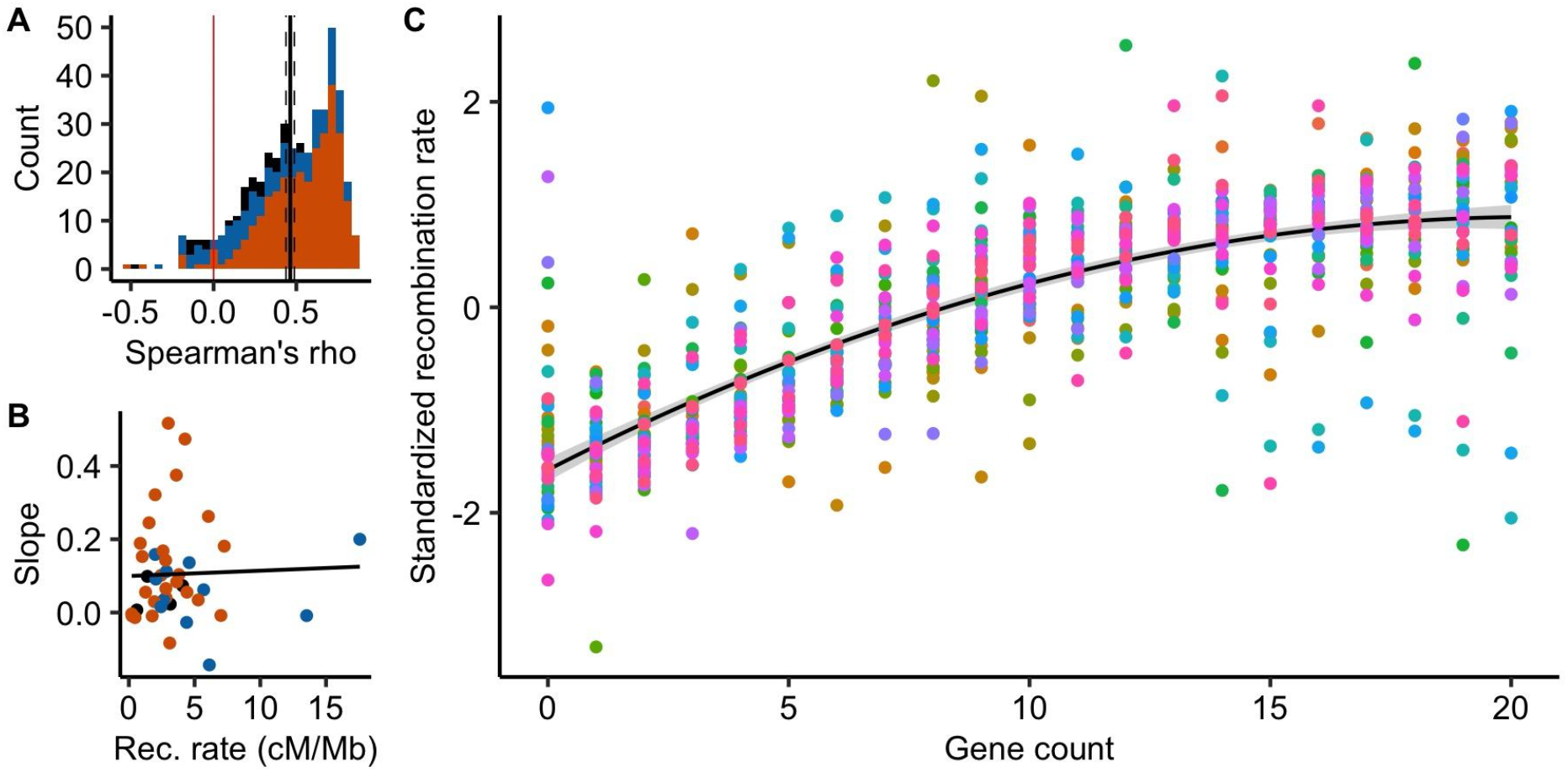
Recombination rates are positively correlated to gene density (n = 483 chromosomes, 41 species). (A) Distribution of chromosome Spearman’s rank correlations between the number of genes and the recombination rate in 100 kb windows. The black vertical line is the mean correlation with a 95% confidence interval (dashed lines) estimated by 1,000 bootstrap replicates. Colours correspond to CO patterns (orange = distal, blue = sub-distal, black = exception). (B) Slopes of the species linear regression between gene count and recombination rates are independent of the species averaged recombination rate (Linear Model, adjusted R^2^ = −0.02, p = 0.83). (C) Standardized recombination rates for each number of genes in a 100 kb window (centred-reduced, chromosomes pooled per species, one colour per species) estimated by 1,000 bootstraps and standardized within species. The gene count was estimated by counting the number of gene starting positions within each 100 kb window. The black line with a grey ribbon is the quadratic regression estimated by linear regression with a 95% parametric confidence interval (Linear Model, adjusted R^2^ = 0.62, p < 0.001).

As for recombination patterns, we classified patterns of gene density along chromosomes in three categories: distal, sub-distal and exceptions (Figure S6). Most species (30 out of 41) were classified in the same gene density and recombination pattern (Table S11). Moreover, when we classified species as a function of recombination patterns, we qualitatively observed the same pattern for gene density and recombination (Figure 8), suggesting that recombination and gene density share the same non-random distribution along the genome.

**Figure 8.**
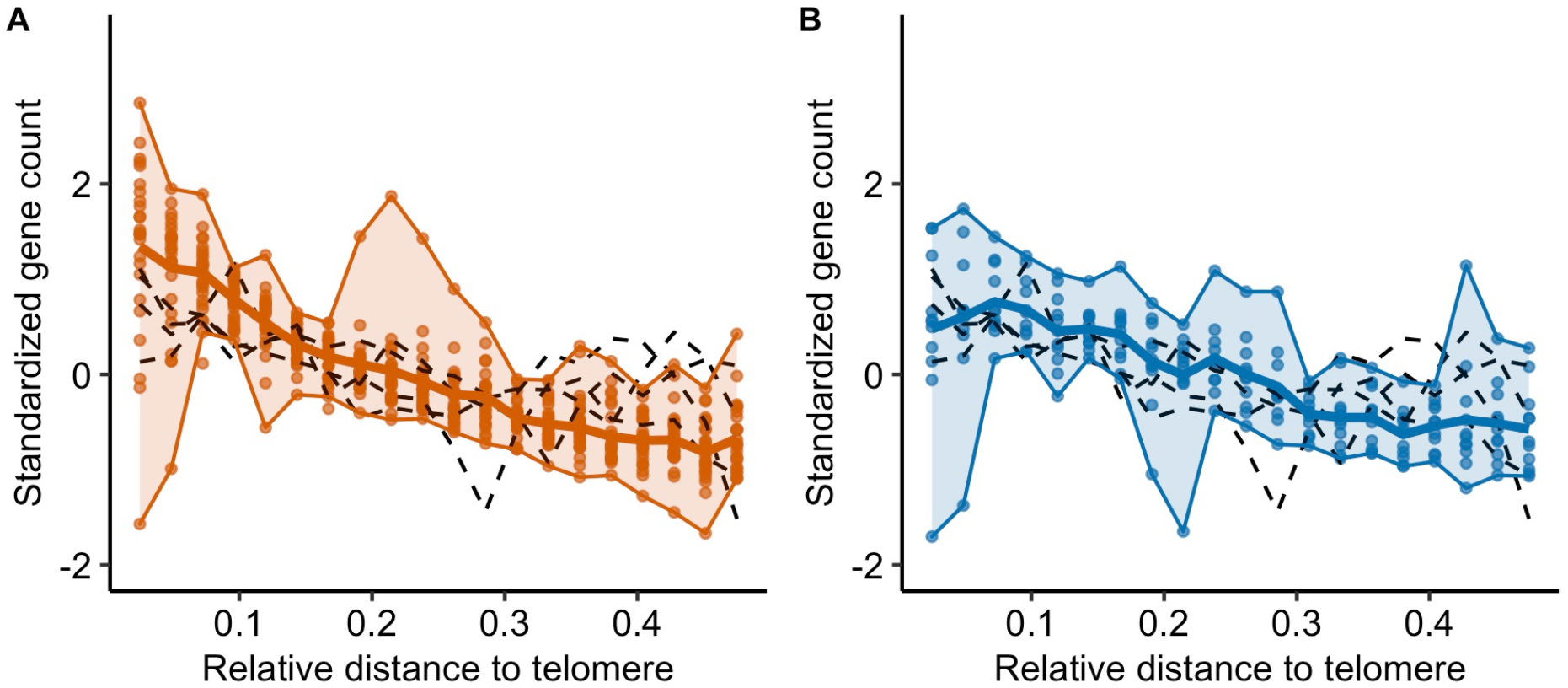
Gene counts patterns along the chromosome are correlated to CO patterns (n = 41 species). Standardized gene count (centred-reduced) as a function of the relative distance from the tip to the middle of the chromosome (genomic distances distributed in 20 bins). We used the same groups as identified for the CO pattern in Figure 6 (distal vs sub-distal) and observed the same patterns along the chromosome. The solid line represents the mean gene count estimated in a bin and the upper and lower boundaries of the ribbon represent the maximum and minimum values in a bin. Patterns that were not classified (4 species with a gene annotation) were represented by loess regression in grey dashed lines. To estimate gene counts in bins of relative distances, chromosomes were split in half, where a distance of 0.5 is the centre of the chromosome. Chromosomes were pooled per species (n = 483 chromosomes).

### Genetic shuffling

We showed that recombination is unevenly distributed in genomes, which should affect how genetic variation is shuffled during meiosis. To quantify how much the genetic shuffling depends on the distribution of COs, we estimated the intrachromosomal component of the genetic shuffling 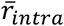 described in equation 10 provided by Veller et al. (2019). The 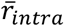 gives a measure of the probability of a random pair of loci to be shuffled by a crossover. As expected, genetic shuffling was positively and significantly correlated with linkage map length (Linear Mixed Model, marginal R^2^ = 0.43, conditional R^2^ = 0.88, p < 0.001, Figure S7). COs clustered in distal regions are supposedly less efficient than COs evenly distributed in the chromosome. At a chromosomal level, linear mixed regression (controlling for a species effect) revealed a low but significant negative effect of the periphery-bias ratio on the genetic shuffling, consistent among species (marginal R^2^ = 0.05, conditional R^2^ = 0.68, p < 0.001, Figure S8). The more COs are clustered in the tips of the chromosome, the lower the chromosomal genetic shuffling. These results verify the analytical predictions of Veller et al. (2019), although the strength of this effect remains weak.

However, the distributions of COs and genes are both non-random and often correlated (Figure S9). Genomic distances measured in base pairs may not be the most appropriate measure of genetic shuffling among functional genomic components. Thus, we measured genomic distances in gene distances (i.e. the cumulative number of genes along the chromosome) instead of base pairs. Marey maps most often appeared more homogeneous when scaled on gene distances instead of base pair distances, with 70% (316 over 450) of Marey maps showing a smaller scale departure from a random distribution (Figures 9, S10, Table S11). Globally, a subset of 30 species have more homogeneous Marey maps with gene distances whereas 11 others are quantitatively more heterogeneous (notably *Capsella rubella* and *Arabidopsis thaliana*), although this could be due to low quality annotations making it difficult to precisely estimate the gene distances for some of them (e.g. *Sesamum indicum*). In most cases, genetic shuffling were slightly higher when gene distances were used instead of base pairs (Figure 10; mean = 0.22 for base pairs; mean = 0.26 for gene distances; Wilcoxon rank sum test with continuity correction, p < 0.001) with a distribution slightly skewed towards higher values for gene distances. Interestingly, the increase in genetic shuffling when scaled with gene distances was more pronounced for longer chromosomes — which are often the most heterogeneous ones, characterized by a distal pattern — whereas we saw little effect on smaller chromosomes characterized by a sub-distal pattern (Linear Mixed Model, marginal R^2^ = 0.16, conditional R^2^ = 0.83, p < 0.001, Figure 10).

**Figure 9.**
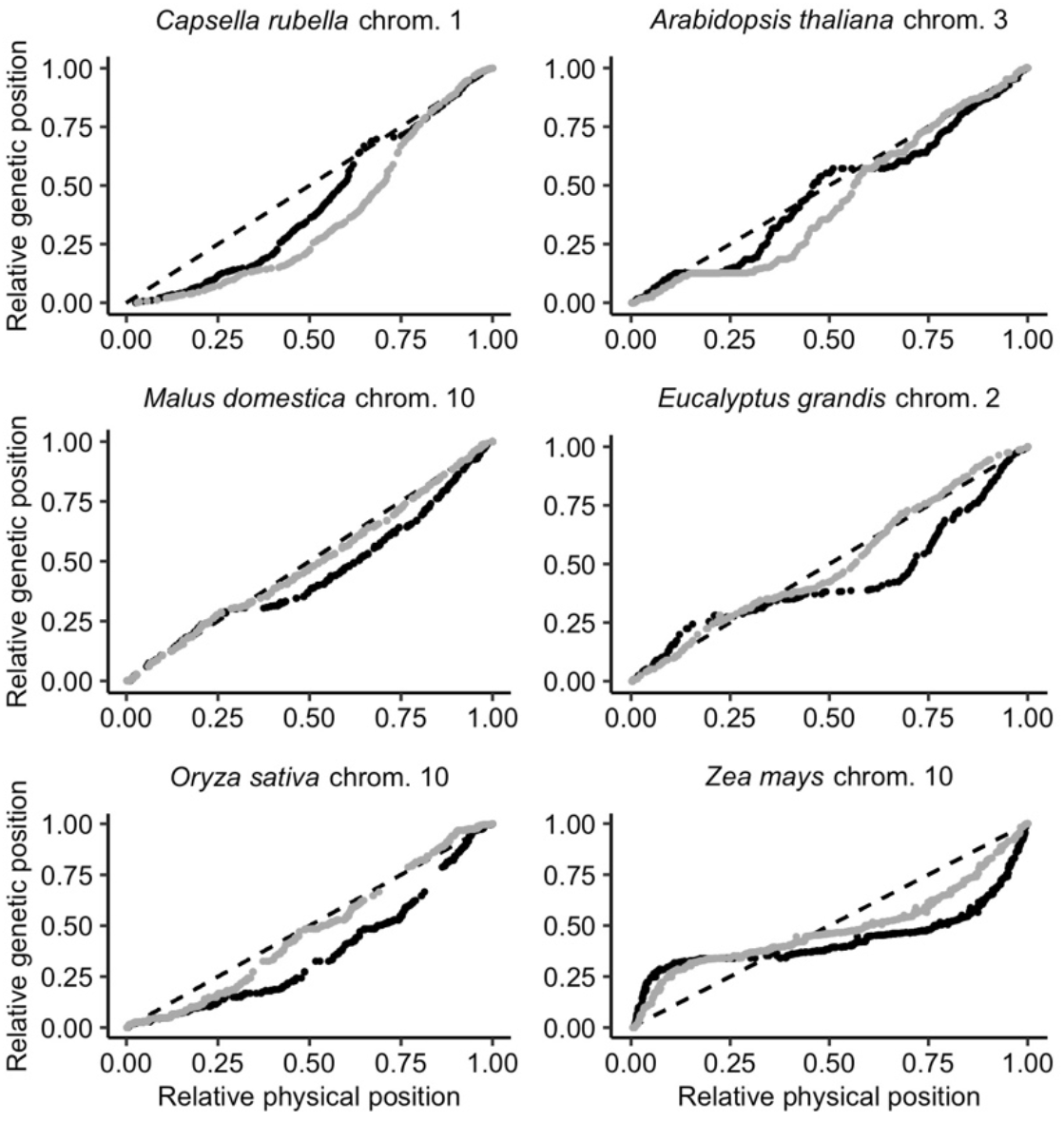
Marey maps of six chromosomes with the relative physical distance expressed in genomic distances (black dots, position in the genome in Mb) or in gene distances (grey dots, position measured as the cumulative number of genes along the chromosome). Marey maps are ordered by ascending chromosome size (Mb). The relative genetic position is the position of the marker on the linkage map. The diagonal dashed line represents a theoretical random distribution of COs along the chromosome.

**Figure 10.**
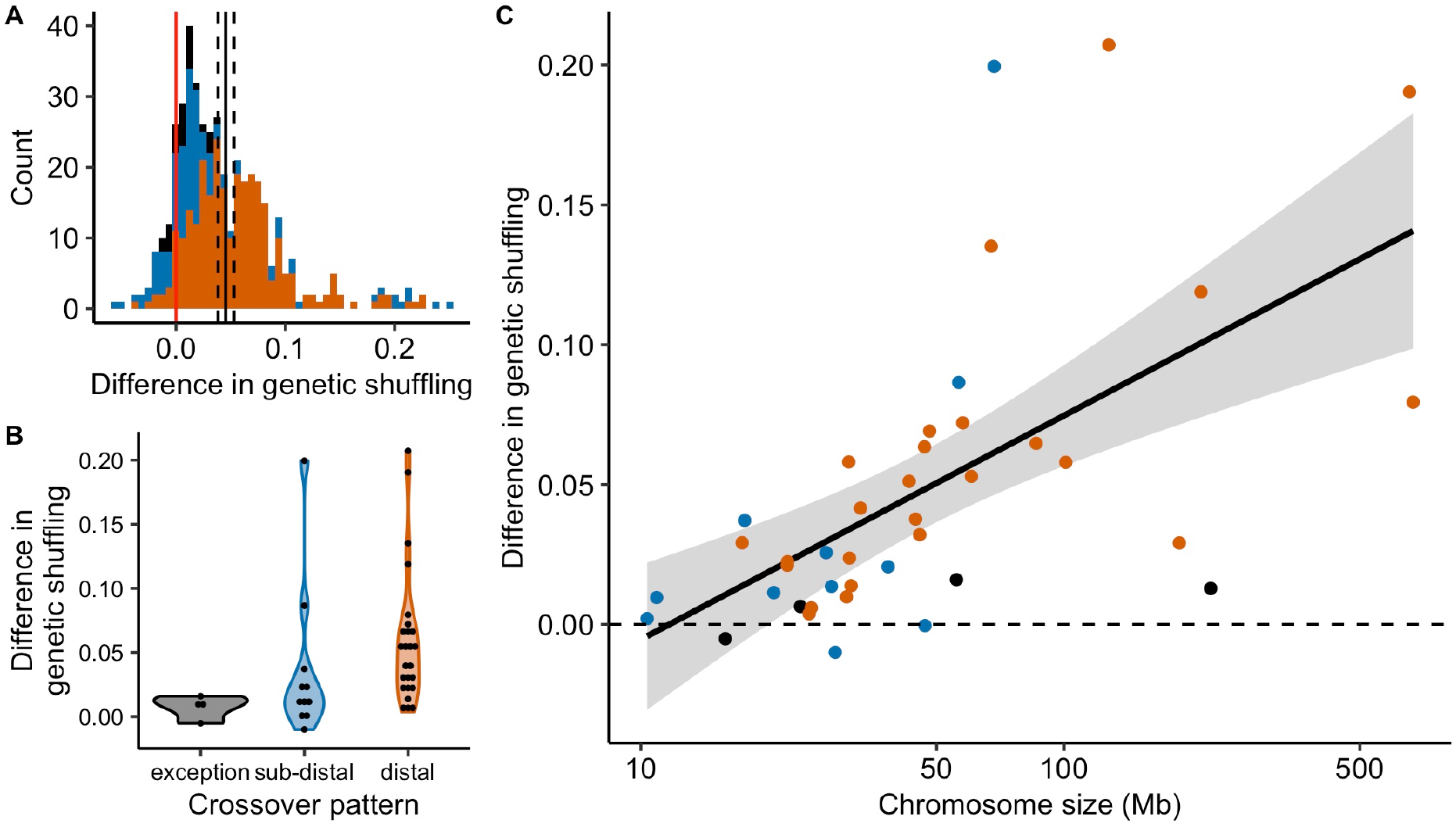
Differences in genetic shuffling between estimates based on genomic distances (Mb) and gene distances (cumulative number of genes). The difference is the genetic shuffling (gene distances) minus the genetic shuffling (genomic distances), thus positive values indicate an increase in the genetic shuffling based on gene distances compared to genomic distances. Colours correspond to CO patterns (orange = distal, blue = sub-distal, black = exception). (A) Distribution of the chromosome differences in the genetic shuffling (n = 444 chromosomes). (B) Distributions of the species difference in the genetic shuffling (n = 41 species, chromosomes pooled). (C) Species differences in the genetic shuffling are positively correlated to the averaged chromosome size (Linear Model, adjusted R^2^ = 0.20, p = 0.002, n = 41, 95% parametric confidence interval).

## Discussion

Based on a large, curated dataset, we provided here, to the best of our knowledge, the largest description of recombination landscapes among flowering plants. In addition to confirming that both the chromosome-wide recombination rate and the heterogeneity of recombination landscapes vary according to chromosome length, we identified two distinct CO patterns and we proposed a new model that builds on the strict telomere model recently suggested by Haenel et al. (2018). Moreover, the consistent correlation between recombination and gene density that we observed suggests that crossover initiation in gene regulatory sequences could be shared among angiosperms. This sheds new light on the efficacy of genetic shuffling and the evolution of recombination landscapes.

### Chromosome size and recombination rate

We showed that, for most species, the smallest chromosome had roughly one or two COs, independently of chromosome size. This is in agreement with the idea that CO assurance is a ubiquitous regulation process among angiosperms (Pazhayam et al., 2021). Moreover, it seems that this constraint imposes a kind of basal recombination rate for each species, on the order of 50/*Sc* cM/Mb, where *Sc* is the size of the lowest chromosome in Mb. Regardless of the genome size (which ranges three orders of magnitude or more), the number of COs remains relatively stable amongst species, most probably under the joint influence of CO assurance, interference and homeostasis (Otto and Payseur, 2019; Stapley et al., 2017; Wang et al., 2015). As a result, averaged recombination rates are negatively correlated to chromosome lengths, as already known in plants (Haenel et al., 2018; Tiley and Burleigh, 2015) and in contrast to fungi and animals (Stapley et al., 2017).

Surprisingly, we found that, within a species, relative chromosome size was a stronger determinant of the genetic map than absolute chromosome size. This suggests that CO interference is proportional to the relative size of the chromosome, as has been empirically observed in some plants (Ferreira et al., 2021). Although it is not clear yet which interference distance unit is the most relevant, genomic distances (in Mb) are excluded in most models of interference in favour of genetic distances (cM) (Foss et al., 1993) or, more likely, the length of the synaptonemal complex in micrometres (Capilla-Pérez et al., 2021; Kleckner et al., 2004; Lloyd and Jenczewski, 2019; Zickler and Kleckner, 2015). Both scales (in genetic distances or in size) of the synaptonemal complex, in micrometres, match our observation of a relative size effect. Within species, genetic maps increase with chromosome size, but among species they are uncorrelated and far less variable than genome sizes, which makes the relative chromosome size the main determinant of recombination rate variations among species. Similarly, physical sizes (in micrometres) at meiosis do not seem to scale with genome size, as chromosomal organization (nucleosomes, chromatin loops) strongly reduces the variation that could be expected given the genome size (Otto and Payseur, 2019).

### Recombination patterns along chromosomes

We observed a global trend towards higher recombination rates in sub-distal regions (Gaut et al., 2007; Haenel et al., 2018). The distal bias increased with chromosome length, in agreement with the conclusions of Haenel et al. (2018), although our methods differ in resolution. We analysed species and chromosomes separately whereas Haenel et al. (2018) used averages over the different patterns, thereby masking chromosome- and species-specific particularities. For example, they did not detect the sub-distal pattern and unclassified exceptions, whereas they seem common among species (14 and 7 species respectively). So far, little is known about the mechanisms that could explain the link between the distal bias and chromosome length. Even if models of CO interference yield similar patterns (Falque et al., 2007; Zhang et al., 2014), the conceptual model of Haenel et al. (2018) is still the only one to explicitly consider chromosome length. The telomere effect is thought to act at a broad chromosome scale over long genomic distances. The decision of double strand breaks (DSBs) to engage in the CO pathway is made early on during meiosis and the early association of telomeres is thought to favour distal COs (Bishop and Zickler, 2004; Higgins et al., 2012; Hinch et al., 2019). In barley, when the relative timing of the first stages of the meiotic program was shortened, COs were redistributed towards proximal regions (Higgins et al., 2012), as later observed in wheat (Osman et al., 2021).

Haenel et al. (2018) proposed that distance to the telomere is driving CO positioning, and therefore it should produce a symmetrical U-shaped pattern along chromosomes. However, a formal test showed that this model was too simple and that centromeres also played a role in the distribution of COs between chromosome arms. The best model (M3: ‘telomere + centromere + one CO per chromosome’) that we have proposed in this work suggests that centromeres do not only have just a local effect but they also influence the symmetry of recombination landscapes over long distances. The local suppression of COs in centromeric regions is well known and largely conserved among species and seems a strong constitutive feature restricted to a short centromeric region, basically the kinetochore (Ellermeier et al., 2010; Fernandes et al., 2019). But the extent of a larger pericentromeric region varies drastically, most probably under the influence of DNA methylation, chromatin accessibility or RNA interference (Choi et al., 2018; Ellermeier et al., 2010; Hartmann et al., 2019; Pan et al., 2011). However, how centromeres may affect CO distribution at larger scales still needs to be determined.

### Diversity of patterns among species

In addition to the role of centromeres, we also observed a departure from the prediction that recombination rates should decrease with the distance to the tip of the chromosome, showing that the distal model is not generally found among plants. We observed at least two different crossover patterns among plant species. Only 34 out of 57 species support a process starting in the tips (distal model), and 16 present the highest recombination rates in sub-distal regions, while seven species remain unclassified, which is at the limit of our visual classification. Globally, the distal pattern and distal bias seem to occur more often in larger chromosomes, but our data lack species with giant genomes. Giant genomes are not rare in plants, and we cannot extrapolate our conclusions to the upper range of the genome size variation (Pellicer et al., 2018). Astonishingly, a low-density genetic map in *Allium* showed higher recombination rates in the proximal regions, which is opposite to the major trend we found (Khrustaleva et al., 2005). Genera with giant genomes such as *Lilium* or *Allium* would have been valuable assets in our dataset, but the actual genomic and linkage data are relatively incomplete (Jo et al., 2017; Shahin et al., 2011).

Contrary to the single pattern described by Haenel et al. (2018), this pattern diversity is more in agreement with what is known of the timing of meiosis and heterochiasmy. Despite the strong conservation of the main meiotic mechanism in plants, differences in the balance between key components may produce distinct CO patterns (de Massy, 2013; Higgins et al., 2012; Kuo et al., 2021; Zelkowski et al., 2019). For example, the ZYP1 telomere-led recombination and the ASY1 protein are two antagonistic forces acting on the formation of the synaptonemal complex in plants (Lambing et al., 2020). In barley and wheat, linearization of the chromosome axis is gradual along the chromosome, initiated in distal regions, whereas early DSBs form in the telomere bouquet (Higgins et al., 2012; Osman et al., 2021). In contrast, chromosome axes are formed at a similar time in *Arabidopsis thaliana* and chromosomes are gradually enriched in ASY1 from the telomeres to the centromeres; a gene-dosage component favours synapsis and ultimately COs towards the proximal regions (Lambing et al., 2020). It appears that the timing of the meiotic programme is important for the distal bias, as it involves changes in the relative contribution of each meiotic component that could explain the re-localization of COs (Higgins et al., 2012; Lambing et al., 2020). Therefore, the different patterns we observed may be explained by the different balance and timing of the expression of shared key regulators of CO patterning such as ZYP1 and ASY1 (Kuo et al., 2021). It is interesting to note that this is also true for mechanistic models of interference. The beam-film model is able to fit both CO patterns, regardless if the tips of the chromosomes have an effect on interference or not, i.e. clamping (Zhang et al., 2014). If there is clamping, mechanical stress culminates in the extremities of the chromosome leading to high CO rates at the periphery where it is released first. In contrast, when clamping is limited, mechanical stress is released in the tips of the chromosome and COs occur further from the tips, until a threshold of mechanical stress is reached. The observed sub-distal pattern fits these predictions.

The two patterns of recombination we describe here can also be observed in opposite sexes within the same species, i.e. heterochiasmy (Capilla-Pérez et al., 2021; Sardell and Kirkpatrick, 2019). Marked heterochiasmy variations between species could influence the resulting sex-averaged recombination landscape (Sardell and Kirkpatrick, 2019). The sex-averaged telomere effect can be thought of as the product of two independent sex-specific landscapes although it is not clear how sex-specific maps ultimately contribute to the sex-averaged one (Johnston et al., 2016; Lenormand et al., 2016). Recombination is usually biased towards the tips of the chromosome in male recombination maps, but is evenly distributed in female maps in most plants (Sardell and Kirkpatrick, 2019). In *Arabidopsis thaliana*, male meiosis has higher CO rates within the tips of the chromosome, as has been observed in other species with large chromosomes, whereas female meiosis is more homogeneously distributed, with the lowest rates found in the distal regions (Capilla-Pérez et al., 2021). Shorter chromosome axes in *A. thaliana* female meiosis could induce fewer DSBs and class II non-interfering COs (Lloyd and Jenczewski, 2019). Conversely, in maize, the distal bias is similar in both sexes, despite higher CO rates for females (Kianian et al., 2018). Heterochiasmy is not universal in plants (Melamed-Bessudo et al., 2016), and we suggest that the variation in recombination landscapes could also result from variation in heterochiasmy among species, as it has been suggested for broad-scale differences in recombination landscapes between *A. thaliana* and its relative *A. arenosa* (Dukić and Bomblies, 2022). This hypothesis should be tested further as more sex-specific genetic maps become available.

### Recombination landscapes, gene density and genetic shuffling

We observed a strong convergence between CO patterns and gene density patterns. This correlation is consistent in our dataset despite possible errors in genome annotation and we also observed two different gene density patterns globally corresponding to similar CO patterns, emphasizing the close link between recombination and gene density. Interestingly, gene density also had a strong correlation with the recombination rate in species with atypical chromosomes. For example, *Camellia sinensis* and *Nelumbo nucifera* have large genomes with homogenous recombination landscapes, whereas large genomes are usually associated with a distal pattern. A recent annotation of a new genome assembly of *Nelumbo nucifera* showed that genes are also evenly distributed along chromosomes at a broad scale (Shi et al., 2020), similar to *Camellia sinensis* (Wei et al., 2018). In wheat and rye, the analysis of the effect of chromosome rearrangement on recombination also suggests that CO localization is more locus-specific than location-specific: after inversions of distal and interstitial segments, COs were relocated to the new position on the distal segment (Lukaszewski, 2008; Lukaszewski et al., 2012). Overall, the parallel between gene density and recombination landscapes, confirmed by these two exceptions, is in agreement with the preferential occurrence of COs in gene regulatory sequences (Choi et al., 2018; He et al., 2017; Marand et al., 2019), and suggests that this may be a general pattern shared among angiosperms. Thus, gene distribution along chromosomes could be a main driver of recombination landscapes simply by determining where COs may preferentially occur. It should be noted that since the gene number is usually positively correlated with chromosome size within a species but is roughly independent of genome size among species, this hypothesis also matches with the relative-size effect discussed above.

However, gene density and recombination rates are both correlated to many other genomic features, such as transposable elements (Marand et al., 2019). The accumulation of transposable elements in low recombining regions would progressively decrease gene density in the region, and would eventually result in a positive correlation between gene density and recombination (Kent et al., 2017). However, the correlation of recombination rates with transposable elements is not always clear and different TE families have opposite correlations (Kent et al., 2017; Underwood and Choi, 2019). The positive association of COs and gene regulatory sequences, including fine-scale correlations, appears more robust (Choi et al., 2013; He et al., 2017; Marand et al., 2019), but causality mechanisms of these multiple interactions still need to be clarified.

Irrespective of the underlying mechanism, our finding implies that the CO distribution ultimately scales with the gene distribution. Therefore, in most species, COs have a more even distribution between genes than between genomic locations (Figure 9). The redistribution of COs towards functional regions could be a simple consequence of COs occurring within gene regulatory sequences, but it has important evolutionary implications such as increasing the genetic shuffling and homogenizing the probability of two random genes to recombine, especially for large genomes that exhibit the strongest difference in genetic shuffling between genes and between genomic locations (Figure 10). Therefore, CO patterning (and not only the global CO rate) could be under selection not only for its direct effect on the functioning of meiosis but also for its indirect effects on selection efficacy (Otto and Payseur, 2019). Recombination decreases linkage disequilibrium and negative interferences between adjacent loci (e.g. Hill-Robertson Interference), and thus locally increases the efficacy of selection. Functional sites are targets for selection (Nachman and Payseur, 2012) and we found higher recombination rates in functional regions, meaning that only a few genes are ultimately excluded from the benefits of recombination, even under the most pronounced distal bias.

Higher recombination rates in gene-rich regions could provide a satisfying explanation as to why the distal bias is maintained among species despite its theoretical lack of efficacy for genetic shuffling (Veller et al., 2019). The association between CO hotspots and gene regulatory sequences is mechanistically driven by chromatin accessibility, but it does not exclude the evolution of the mechanism itself towards the benefits of recombining more in gene-rich regions (Lenormand et al., 2016). However, slight variations in genetic shuffling caused by the non-random distribution of COs are less likely to be under strong selection compared to stabilizing selection on molecular constraints for chromosome pairing and segregation (Ritz et al., 2017), although interference is sometimes likely to evolve towards relaxed physical constraints (Otto and Payseur, 2019). In addition, the intra-chromosomal component of the genetic shuffling is a small contributor to the genome-wide shuffling rate, as a major part is due to independent assortment among chromosomes (Veller et al., 2019) even though there may be significant selective pressure towards more recombination between genes within chromosomes. Our estimates for the chromosomal genetic shuffling do not reach the theoretical optimal value of 0.5. The pattern is not absolute, and a fraction of genes remains in low recombining regions. In grass species, up to 30% of genes are found in recombination deserts and are not subject to efficient selection (e.g. Mayer et al., 2011). Finally, it is still an open question as to whether this global distribution of COs in gene regulatory sequences is advantageous for the genetic diversity and adaptive potential of a species (Pan et al., 2016).

### Conclusion

Our comparative study only demonstrates correlations, and not mechanisms, but helps to understand the diversity and determinants of recombination landscapes in flowering plants. Our results partly confirm previous studies based on fewer species (Haenel et al., 2018; Stapley et al., 2017; Tiley and Burleigh, 2015) while bringing new insights that alter previous conclusions thanks to a detailed analysis at the species and chromosome levels. Two main and distinct CO patterns emerge across a large set of flowering plant species; it seems likely that chromosome structure (length, centromere) and gene densities are the major drivers of these patterns, and the interactions between them raise questions about the evolution of complex genomic patterns at the chromosome scale (Gaut et al., 2007; Nam and Ellegren, 2012). The new large and curated dataset we provide in the present work should be useful for addressing such questions and testing future evolutionary hypotheses regarding the role of recombination in genome architecture.

## Materials and Methods

### Data preparation

To build recombination maps, we combined genetic and genomic maps in angiosperms that had already been published in the literature. We conducted a literature search to collect sex-averaged genetic maps estimated on pedigree data – with markers positions in centiMorgans (cM). The keywords used were ‘genetic map’, ‘linkage map’, ‘genome assembly’, ‘plants’ and ‘angiosperms’, combined with ‘high-density’ or ‘saturated’ in order to target genetic maps with a large number of markers and progenies. Additionally, we carried out searches within public genomic databases to find publicly available genetic maps. Only species with a reference genome assembly at a chromosome level were included in our study (a complete list of genetic maps with the associated metadata is given in Tables S1, S2). As much as possible, genomic positions along the chromosome (Mb) were estimated by blasting marker sequences on the most recent genome assembly. Genome assemblies with annotation files at a chromosome-scale were downloaded from NCBI (https://www.ncbi.nlm.nih.gov/) or public databases. Marker sequences were blasted with ‘blastn’ and a 90% identity cutoff. Markers were anchored to the genomic position of the best hit. When the sequence was a pair of primers, the mapped genomic position was the best hit between pairs of positions showing a short distance between the forward and reverse primer (< 200 bp). In a few exceptions (see Table S1), genomic positions were mapped on a close congeneric species genome and the genomic map was kept if there was good collinearity between the genetic and genomic positions. Chromosomes were numbered as per the reference genome assembly. When marker sequences were not available, we kept the genomic positions published with the genetic map. The centromere position was retrieved from the literature (i.e. the centromeric index, the ratio of the short arm length versus the total chromosome length). The total genomic length was estimated by the length of the chromosome sequence in the genome assembly. The total genetic length was corrected using Hall and Willis’s method (Hall and Willis, 2005) which accounts for undetected events of recombination in distal regions by adding *2s* to the length of each linkage group (where *s* is the average marker spacing in the group).

We selected genetic and genomic maps after stringent filtering and corrections. We assumed that markers must follow a monotone increasing function when plotting genetic distances as a function of genomic distances in a chromosome (i.e. the Marey map) to guarantee collinearity between the genetic map and the reference genome. If necessary, genetic maps were reoriented so that the Marey map function is increasing. In a first step, Marey maps with fewer than 50 markers per chromosome were removed, although a few exceptions were visually validated (maps with ~30 markers). Marey maps with more than 10% of the total genomic map length missing at one end of the chromosome were removed. Marey maps with obvious artefacts and assembly mismatches (e.g. large inversions, large gaps) were removed. Markers clearly outside the global trend of the Marey map were visually filtered out. The Marey map approach is a graphical method, so figures were systematically produced at each step as a way to evaluate the results of the filtering and corrections. Finally, when multiple datasets were available for the same species, we selected the dataset with the highest marker density – in addition to visual validation – to maintain a balanced sampling and avoid pseudo-replicates of the same chromosome.

### Estimates of local recombination rates

Local recombination rates along the chromosome were estimated with the Marey map approach described in the MareyMap R package (Rezvoy et al., 2007). The mathematical function of the Marey map was interpolated with a two-degree polynomial loess regression. Each span smoothing parameter was calibrated by 1,000 iterations of hold-out partitioning (random sampling of markers between two subsets; 2/3 for training and 1/3 for testing) with the Mean Squared Error of the loess regression as a goodness-of-fit criterion. The possible span ranged from 0.2 to 0.5 and was visually adjusted for certain maps. The local recombination rate was the derivative of the interpolated smoothed function in fixed 100 kb and 1 Mb non-overlapping windows. Negative estimates were not possible as we assumed a monotonously increasing function and negative recombination rates were set to zero. The 95% confidence intervals of the recombination rates were estimated by 1,000 bootstrap replicates of the markers. The quality of the estimates was checked using the correlation between the 100 kb and 1 Mb windows.

### The broken stick model

The spatial structure of recombination landscapes across species and chromosomes is a major feature of recombination landscapes. Applied to the distribution of recombination in Marey maps, our implementation of the broken stick model seemed effective to visualize the broad-scale variation of recombination rates (White and Hill, 2020). We divided the Marey map in *k* segments of equal genetic size and then calculated the relative genomic size of each segment. Under the null model (i.e. random recombination), one expects *k* segments of equal genomic size 1/*k*. The relative recombination rate was estimated by the log-ratio of the relative genetic size divided by the relative genomic size. Given the observation that most recombination landscapes are broken down into at least three segments (White and Hill, 2020), we arbitrarily chose a number of segments *k* = 10 to reach a good resolution (a larger *k* did not show any qualitative differences).

### Crossover patterns and the periphery-bias ratio

We investigated the spatial bias towards distal regions of the chromosome in the distribution of recombination by estimating recombination rates as a function of relative distances to the telomere (i.e. distance to the nearest chromosome end). Chromosomes were split by their midpoint and only one side was randomly sampled for each chromosome to avoid pseudo-replicates and the averaging of two potentially contrasting patterns on opposite arms. The relative distance to the telomere was the distance to the telomere divided by total chromosome size, then divided into 20 bins of equal relative distances. A periphery-bias ratio metric similar to the one presented in Haenel et al. (2018) was estimated to measure the strength of the distal bias. We divided the recombination rates in the tip of the chromosome (10% on each side of the chromosome, and one randomly sampled tip) by the mean recombination rate of the whole chromosome. We investigated the sensitivity of this periphery-bias ratio to the sampling scale by calculating the ratio for many distal region sizes (Figure S11).

### Testing centromere or telomere effects

We searched the literature for centromeric indices (ratio of the short arm length divided by the total chromosome length) established by cytological measures. When we had no information about the correct orientation of the chromosome (short arm/long arm), the centromeric index was oriented to match the region with the lowest recombination rate of the whole chromosome (i.e. putative centromere). To determine if telomeres and centromeres play a significant role in CO patterning, we fitted empirical CO distributions to three theoretical models of CO distribution. In the following equations, *d(x)* is the relative genetic distance at the relative genomic position *x*, and *a* is a coefficient corresponding to the excess of COs per genomic distance. Under the strict ‘telomere’ model (1), we assumed that only telomeres played a role in CO distribution, i.e. an equal distribution of COs on both sides of the chromosome (i.e. *d*(1/2) = *d*(1) – *d*(1/2), such that 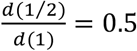. The ‘telomere + centromere + one mandatory CO per arm’ model (2) assumed at least one CO per chromosome arm and a relative genetic distance of each chromosome arm proportional to its relative genomic size, corresponding to the role of centromere position, denoted *d(c)*. We have *d*(*c*) = 50 + *a* × *c* and *d*(1) – *d*(*c*) = 50 + *a* × (1 – *c*), such that 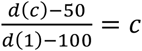. Lastly, the ‘telomere + centromere + one CO per chromosome’ model (3) assumed at least one CO per chromosome and a relative genetic distance within the chromosome proportional to its relative genomic distance. We have *d*(*c*) = *c* × 50 + *a* × *c* and *d*(1) – *d*(*c*) = (1 – *c*) × 50 + *a* × (1 – *c*), such that 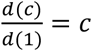. The three competing models were compared with a linear regression between empirical and theoretical values, based on the adjusted R^2^ and AIC-BIC criteria among chromosomes. The number of species supporting each model was calculated based on the adjusted R^2^ within species, for all species with at least five chromosomes.

### Gene density

We retrieved genome annotations (‘gff’ files) for genes, coding sequences and exon positions, preferentially from NCBI and otherwise from public databases (41 species). We estimated gene counts in 100 kb windows for recombination maps by counting the number of genes with a starting position falling inside the window. For each gene count, we estimated the species mean recombination rate and its confidence interval at 95% by 1,000 bootstrap replicates (chromosomes pooled per species). Most species had rarely more than 20 genes over a 100 kb span and variance dramatically increased in the upper range of the gene counts, and therefore we pruned gene counts over 20 for graphical representation and statistical analyses.

### Genetic shuffling

To assess the efficiency of the recombination between chromosomes and species, we calculated the measure of intra-chromosomal genetic shuffling described by Veller et al. (2019). To have even sampling along the chromosome, genetic positions (cM) of 1,000 pseudo-markers evenly distributed along genomic distances (Mb) were interpolated using a loess regression on each Marey map, following the same smoothing and interpolation procedure as for the estimation of the recombination rates. The chromosomal genetic shuffling 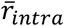 were calculated as per the intra-chromosomal component of the equation 10 presented in Veller et al. (2019). For a single chromosome,

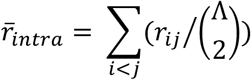

where Λ is the total number of loci, 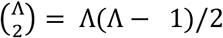 and *r_ij_* is the rate of shuffling for the locus pair *(i, j)*. For the intra-chromosomal component 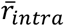, the pairwise shuffling rate was only calculated for linked sites, i.e. loci on the same chromosome. This pairwise shuffling rate was estimated by the recombination fraction between loci *i* and *j*. Recombination fractions were directly calculated from Haldane or Kosambi genetic distances between loci by applying a reverse Haldane function (1) or reverse Kosambi function (2), depending on the mapping function originally used for the given genetic map.

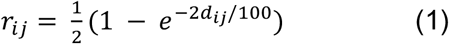

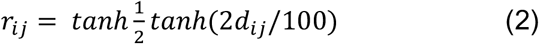

We also estimated marker positions in gene distances instead of genomic distances (Mb) to investigate the influence of the non-random distribution of genes on the recombination landscape. Gene distances were the cumulative number of genes along the chromosome at a given marker’s position. Splicing variants and overlapping genes were counted as a single gene. The genetic shuffling was re-estimated with gene distances instead of genomic distances to consider a genetic shuffling based on the gene distribution, as suggested by Veller et al. (2019). To compare the departure from a random distribution along the chromosome among both types of distances (i.e. genomic and genes), we calculated the Root Mean Square Error (RMSE) of each Marey map and for both distances. To assess if the distribution of genes influenced the heterogeneity of recombination landscapes, the type of distance with the lower RMSE was considered as the more homogeneous landscape. However, this measure for gene distances is sensitive to annotation errors and artefacts. False negatives are therefore expected (when Marey maps were assessed as more homogeneous in genomic distances while the inverse is true) and this classification remains conservative.

### Statistical analyses

All statistical analyses were performed with R version 4.0.4 (R Core Team, 2019). We assessed statistical relationships with the non-parametric Spearman’s rank correlation and regression models. Linear Models were used for regressions with species data since we did not detect a phylogenetic effect. The structure in the chromosome dataset was accounted for by Linear Mixed Models (LMER) implemented in the ‘lme4’ R package (Bates et al., 2015, p. 4) and the phylogenetic structure was tested by fitting the Phylogenetic Generalized Linear Mixed Model (PGLMM) of the ‘phyr’ R package (Ives et al., 2019). The phylogenetic time-calibrated supertree used for the covariance matrix was retrieved from the publicly available phylogeny constructed by Smith and Brown (Smith and Brown, 2018). Marginal and conditional R^2^ values for LMER were estimated with the ‘MuMIn’ R package (Bartoń, 2020). Significance of the model parameters was tested with the ‘lmerTest’ R package (Kuznetsova et al., 2017). We selected the model based on AIC/BIC criteria and diagnostic plots. Reliability and stability of the various models were assessed by checking quantile-quantile plots for the normality of residuals and residuals plotted as a function of fitted values for homoscedasticity. Model quality was checked by the comparison of predicted and observed values. Given the skewed nature of some distributions, we used logarithm (base 10) transformations when appropriate. For comparison between species, statistics were standardized (i.e. by subtracting the mean and dividing by standard deviation). Mean statistics and 95% confidence intervals were estimated by 1,000 bootstrap replicates.

## Notes

### Competing Interest Statement

The authors have declared no competing interest.

